# A novel camera trap design for studying wildlife in mountain glacier ecosystems yields new insight for glacier biodiversity in the Pacific Northwest, USA

**DOI:** 10.1101/2021.12.10.472172

**Authors:** Scott Hotaling, Jordan Boersma, Neil A. Paprocki, Alissa Anderson, Logan Whiles, Lucy Ogburn, Sophia Kasper, Catharine White, Daniel H. Thornton, Peter Wimberger

**Author notes:** **Correspondence:** Scott Hotaling, Department of Watershed Sciences, Utah State University, Logan, UT, USA.

## Abstract

**Context:** The global recession of glaciers and perennial snowfields is reshaping mountain ecosystems. Beyond physical changes to the landscape and altered downstream hydrology, the implications of glacier decline for biodiversity are poorly known. Before predictions can be made about how climate change will affect wildlife in glacier-associated ecosystems, a more thorough accounting of the role that glaciers play in species’ life histories is needed. However, typical approaches for documenting wildlife presence and behavior—remote camera traps—are difficult to use in glaciated terrain due to limited options for securing them (e.g., no trees) and dramatic seasonal changes in snowpack.

**Aims:** In this study, we sought to test a novel camera trap designed for glaciated mountain ecosystems. We also aimed to use this approach to gain insight into wildlife and human usage of a mountain glacier in western North America.

**Methods:** We deployed an elevational transect of uniquely designed camera traps along the western margin of the Paradise Glacier, a rapidly receding mountain glacier on the south side of Mount Rainier, WA, USA. Our simple camera trap design consisted of a wildlife camera attached to a camouflaged cylindrical cooler filled with snow and rocks.

**Key results:** Our camera design proved ideal for a mountain glacier ecosystem and from June to September 2021, we detected at least 16 vertebrate species (seven birds, nine mammals) over 770 trap nights using glacier-associated habitats. Humans, primarily skiers, were the most common species detected, but we also recorded 99 observations of wildlife (birds and mammals). These included three species of conservation concern in Washington: wolverine (*Gulo gulo*), Cascade red fox (*Vulpes vulpes cascadensis*), and white-tailed ptarmigan (*Lagopus leucura*).

**Conclusions:** Collectively, our results provide proof-of-concept for a novel camera trap design that is ideal for treeless, perennially snow-covered landscapes and revealed a rich diversity of wildlife using mountain glacier habitat in the Pacific Northwest. We highlight the global need for similar studies to better understand the true scale of biodiversity that will be impacted by glacier recession in mountain ecosystems.

## Introduction

As climate change proceeds and mountain glaciers are lost, there is a pressing need to understand how the loss of glacier ice will impact habitats and ecosystems (Hotaling *et al*. 2017; Milner *et al*. 2017). Well-known impacts of glacier recession include the loss of key meltwater resources which transport nutrients to downstream habitats (Ren et al. 2019), underpin hydropower infrastructure (Finger *et al*. 2012), and provide key habitats for specialized aquatic communities (Giersch *et al*. 2017; Hotaling *et al*. 2019; Muhlfeld *et al*. 2020). Often overlooked in this discussion, however, is the impact of glacier loss on the biodiversity living on and adjacent to glaciers (Hardy *et al*. 2018; Schano *et al*. 2021; Stibal *et al*. 2020).

The biodiversity of glacier ecosystems has gained appreciation in recent decades. Long considered devoid of life, it is now established that glaciers host a thriving, multi-kingdom community (Anesio *et al*. 2017; Shain *et al*. 2021), particularly in the uppermost layer of snow and ice where intense solar radiation, atmospheric deposition, and available water during melt yield conditions that are more hospitable to life (Anesio and Laybourn-Parry 2012; Hotaling *et al*. 2021). Still, however, most studies have focused on the most dominant life form on glaciers— microbes—leaving larger-bodied residents of glaciers and surrounding habitats overlooked.

Globally, 36 species of birds and mammals have been observed using glacier and perennial snow habitat for some portion of their life history (Rosvold 2016). Mammals typically use these habitats for relief from lower elevation conditions during warmer parts of the year, whereas birds commonly forage on arthropods and seeds that have been atmospherically deposited on the ice (Antor 1995; Hotaling *et al*. 2021; Rosvold 2016). However, closer links between wildlife and glacier habitats have also been described. In western North America, the high abundance of glacier ice worms (*Mesenchytraeus solifugus*)—which are commonly present at densities >200 per m^2^ on glaciers during summer—may be a key resource subsidy for nesting birds (e.g., gray-crowned rosy finches, Hotaling *et al*. 2020). On glaciers of the high Andes in South America, the white-winged diuca finch nests directly within crevasses (Hardy *et al*. 2018). Throughout higher latitudes of the Northern Hemisphere, wolverines cache food in snow and ice, and the persistence of spring snow cover and cache longevity has been hypothesized as a primary reason underpinning their cold, snowy distributions (i.e., the “refrigeration-zone” hypothesis, Inman *et al*. 2012).

While links between larger-bodied vertebrates and icy habitats have been made for a range of species, these observations typically stem from focused species-specific studies and likely underestimate the true scale of wildlife use in the cryosphere (the global collection of Earth’s frozen waters). Beyond researcher focus on specific species, this underestimation likely also stems from two other, intertwined challenges. Mountain ecosystems, and particularly glaciers, are poorly studied relative to more convenient habitats (e.g., those in close proximity to universities and museums, Freitag *et al*. 1998). Moreover, mountain ecosystems have additional challenges associated with their study. Indeed, for mountain research, typical field methods are often difficult to implement due to, for example, the remoteness, rugged terrain, and harsh weather conditions (and high rates of snowfall) associated with these habitats. For instance, most camera trapping studies affix cameras to trees or T-Post stakes driven into soil (Rich *et al*. 2019; Rovero *et al*. 2010). In mountain glacier ecosystems, trees are typically absent, soil is usually covered by deep snowpack or non-existent due to glacier activity scouring the landscape down to bedrock, and affixing cameras to boulders limits locations for deployment and timing (e.g., early season deployments might not be possible due to boulders still being covered by seasonal snow). Thus, an easy to transport, flexible, but robust camera trap design to ameliorate the many challenges of high mountain ecosystems is needed.

In this study, we sought to gain a more holistic understanding of human and wildlife usage of alpine glacier habitat while also testing a simple camera trap designed to alleviate the difficulties associated with high-mountain ecosystems. We deployed our camera-trapping array along an elevational transect of a mountain glacier on Mount Rainier, Washington, USA. While we expected our camera trap array to detect many species known to occur in these ecosystems (e.g., marmot), we also expected to observe species not previously associated with glacier or snow ecosystems. If detected, these results would underscore the likely underestimation of glacier-linked vertebrate biodiversity globally and the pressing need to better understand how climate change may impact enigmatic mountain species.

## Materials and methods

### Study area

Mount Rainier is a large stratovolcano in south-central Washington, USA (Figure 1a), and the centerpiece of Mount Rainier National Park (MORA). At 4,392 m (14,411 feet), Mount Rainier is the highest peak in Washington state and is flanked by 29 named glaciers covering ∼78 km^2^ (Beason 2017). The Paradise Glacier lies on the southern side of the peak—with lower and upper extents at 2,100 and 2,585 m, respectively—and is one of the smaller glaciers in MORA at ∼0.6 km^2^. Like most glaciers around the world, Mount Rainier’s glaciers are in rapid retreat. From 1896 to 2015, glaciated area on Mount Rainier has decreased by ∼40%, a loss of 50.6 km^2^ (Beason 2017). The Paradise Glacier, which was part of a larger Paradise-Stevens Glaciers complex in 1896, has retreated by 87% over the same period (Beason 2017). Still, Mount Rainier and its glaciers remain some of the snowiest places on Earth. Annual snowfall at 1,646 m—as measured near the park visitor center at Paradise (Figure 1)—has averaged 1,620 cm (639 inches) from 1920 - 2020 (National Park Service 2021a). On Mount Rainier, treeline occurs from roughly 1,829 - 1,981 m (6,000 - 6,500 feet).

**Figure 1.**
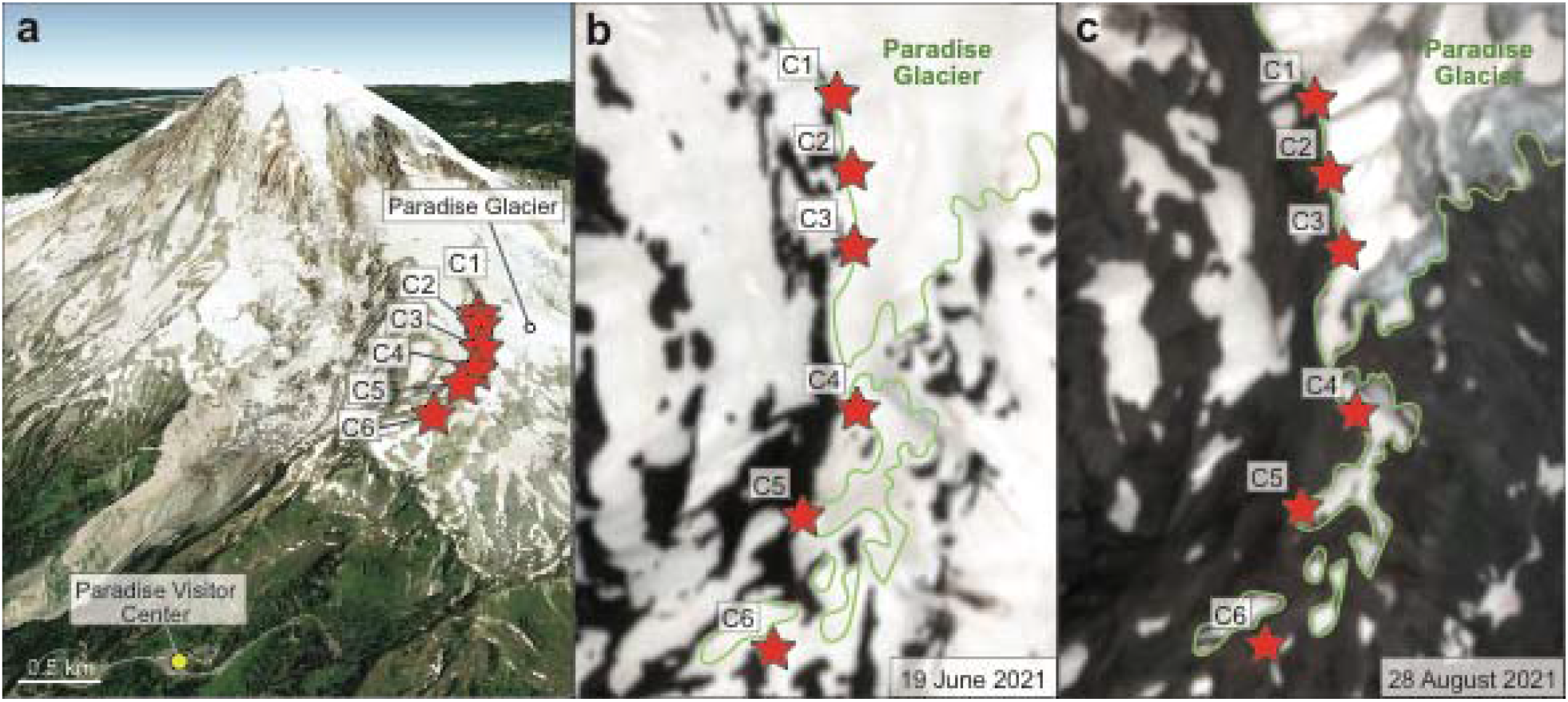
(a) Mount Rainier, Washington, USA, viewed from the south with our camera locations labeled. Satellite imagery from Google Earth. (b, c) Sentinel-2 satellite imagery of our camera locations when they were first deployed (b) and once the bulk of seasonal snow had melted (c). The outline of the Paradise Glacier (thin green line) in (b) and (c) was approximated from Beason (2017).

### Experimental design

From 17 June 2021 to 31 October 2021, we deployed wildlife cameras (Browning Strike Force 850 Extreme, Browning, USA) along the western margin of the Paradise Glacier (Figures 1b,c). However, we only analyzed data through 25 September 2021 because variable but often persistent snowpack either buried or obscured cameras during the final five weeks of the field season (25 September - 31 October 2021). We placed six cameras along an elevational transect from 2,164 to 2,469 m (7,100 - 8,100 feet) and re-visited the cameras every 3-5 weeks to adjust positioning, exchange memory cards, and perform similar routine activities. We focused our cameras to target the margins of the glacier where talus and boulders give way to ice. Because the Paradise Glacier is such a dynamic environment during summer, with a dramatically shifting physical surface (both in area and elevation), we moved the cameras as needed to maintain close proximity (generally 0-10 m) to the ice surface. When triggered, cameras took bursts of five photos in quick succession with a 30-second rest period between triggers. While our camera traps were largely reliable and stayed in working order throughout the study, we did experience a memory card failure at C5 from 20 August - 25 September. Due to the variable terrain and study goals, we made no attempts to control for the amount of area each camera could “see” which likely affected the comparability of our results across cameras.

To account for the challenges of the alpine glacier ecosystem, including the need to move the cameras throughout the melt season, we employed a unique camera trap design (Figures 2a,b). Cameras were mounted to 5-gallon (18.9 liter) cylindrical Igloo^©^ coolers that were covered in camouflage duct tape with small, camouflage ratchet straps. The coolers were filled with snow and loose rocks from the trapping area and sealed with ⅛” coated steel cable and a small U-clamp. Because we expected visitors to notice our cameras despite efforts to blend them into the landscape, we added a small label to the top of each trap noting its purpose with contact information. The low-profile but robust footprint of our camera traps when the coolers were filled with snow/rock provided a stable platform that remained in place despite high winds and an unprecedented heat wave that occurred during the course of our study and rapidly melted a large amount of snowpack (Pelto *et al*. 2022).

**Figure 2.**
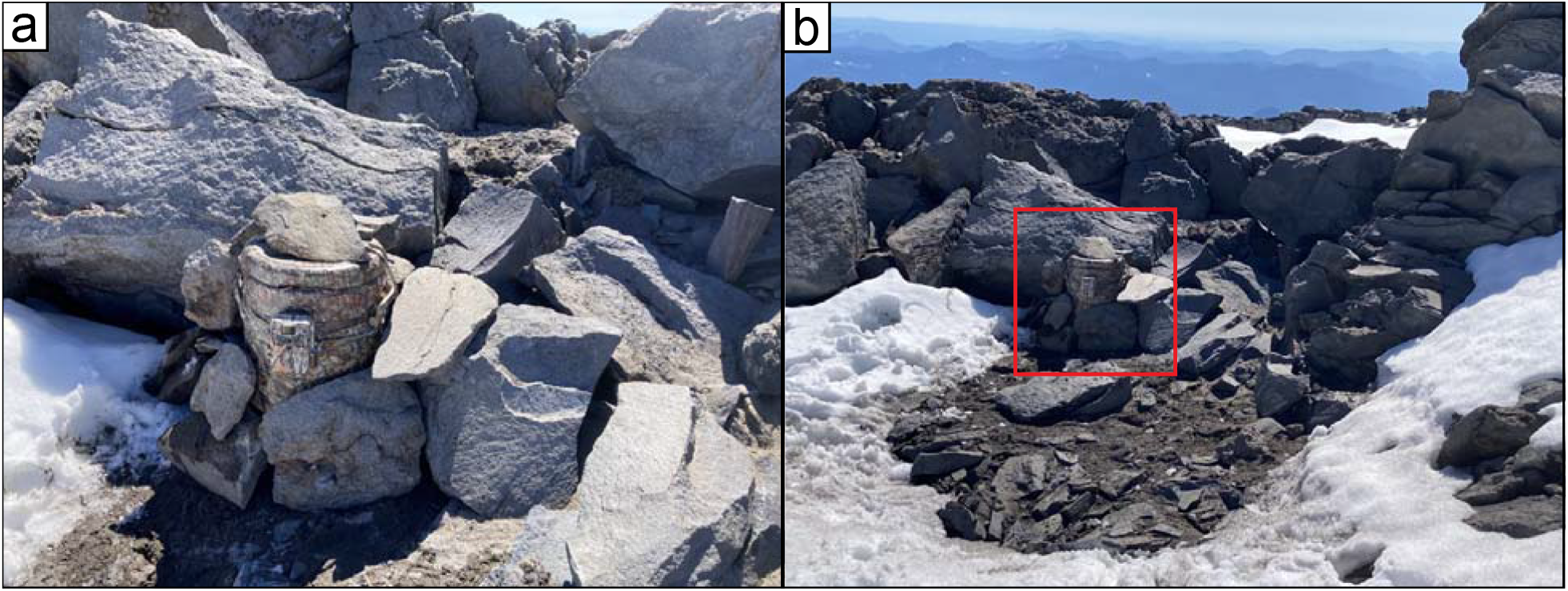
(a) A close-up view of an example camera trap on the margin of the Paradise Glacier on Mount Rainier, Washington, USA. (b) A more distant view of the camera (within the red square) showing how it blends into the surrounding talus. Photographs: © Scott Hotaling.

### Animal identification and data analysis

Images were visually inspected for any evidence of vertebrate detection (see example images in Figure 3). When humans or wildlife (birds and mammals) were detected, they were identified to the lowest taxonomic level possible by consulting guides for Mount Rainier National Park and the Pacific Northwest as well as our own research team’s expertise in mammals (A.A. and L.W., specifically) and birds (J.B., N.A.P., and P.W., specifically).Given the proximity of our cameras to glacier ice or perennial snow, we considered any species recorded in this study to have a link to glacial habitats. This definition is slightly expanded relative to Rosvold (2016), the most similar study to ours, which focused exclusively on birds and mammals that were directly on snow and ice.

**Figure 3.**
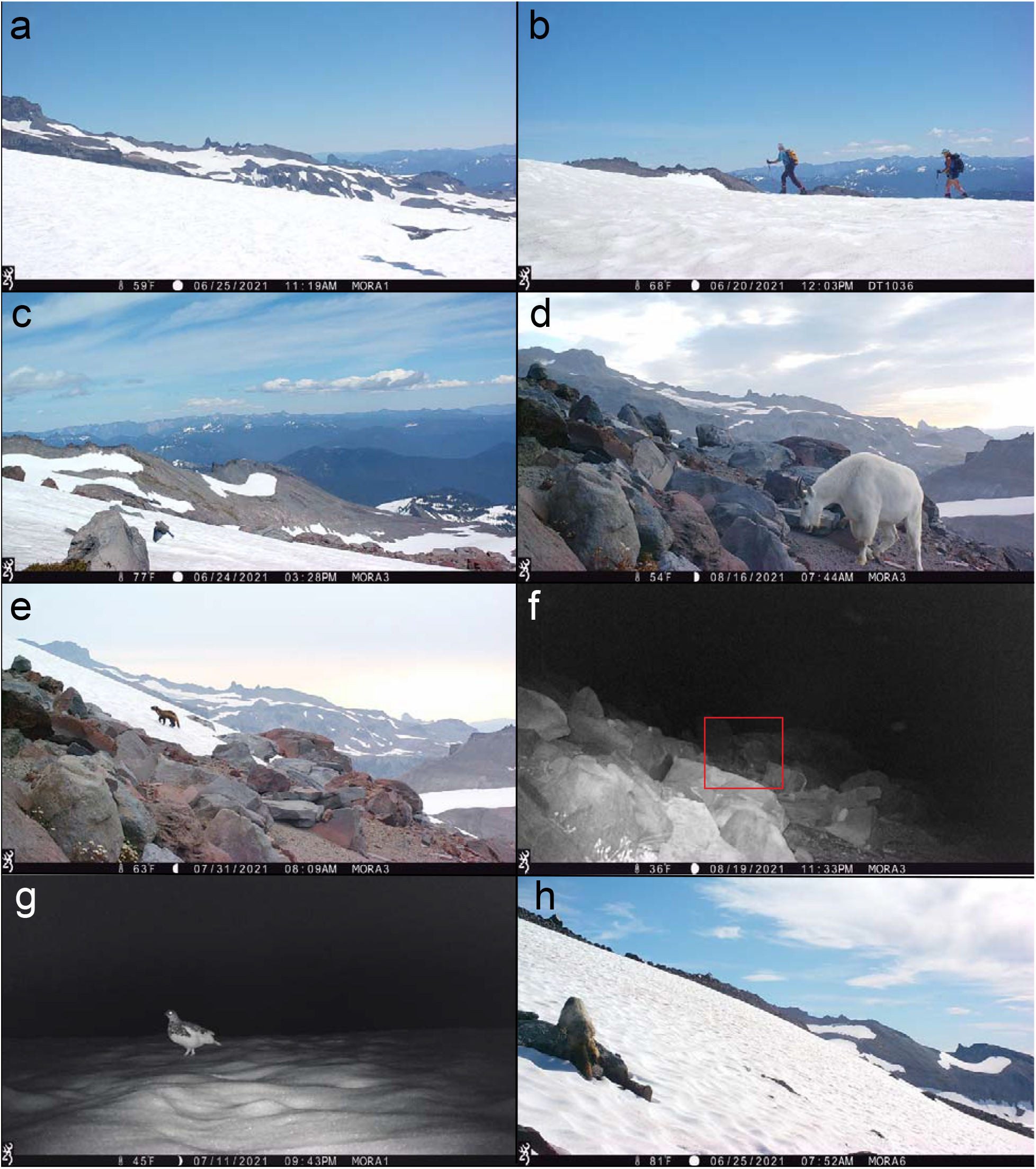
Examples of taxa detected with camera traps along the western margin of the Paradise Glacier, Mount Rainier, WA, USA. (a) Common raven (*Corvus corvax*), (b) humans (*Homo sapiens*), (c) mountain bluebird (*Sialia corrucoides*), (d) mountain goat (*Oreamnos americanus*), (e) wolverine (*Gulo gulo*), (f) Cascade red fox (*Vulpes vulpes cascadensis*), (g) white-tailed ptarmigan (*Lagopus leucura*), (h) hoary marmot (*Marmota caligata*).

When multiple images from a camera burst included the same organism(s), it was only counted once. The same is true for multiple bursts within minutes of each other. However, because our camera array was arranged along an elevational transect, it is possible, and perhaps likely, that some humans or wildlife were detected multiple times across different cameras. We did not attempt to control for multiple identifications in space nor time beyond the methods described above. Photos of the research team were also excluded from all analyses. Observations were totaled and analyzed for each of the 14 weeks of the study. We note that the final “week” (week 14) contained two extra days due to where the study endpoint fell (25 September) relative to the end of that particular week (23 September).

To test for a relationship between human and wildlife occurrences in our data set, we U a series of statistical analyses in R v3.6.3 (R Core Team 2021). First, we grouped all human and wildlife observations separately for each camera on a week-by-week basis. Then, we tested for normality of the human and wildlife data sets using a Shapiro-Wilk normality test (“shapiro.test”). Because both data sets were significantly different from normal (*P*, Shapiro-Wilk < 0.001), we used correlation analyses with a Spearman’s rank correlation. We tested for a correlation between the number of human versus wildlife detections across all cameras and weeks in the study.

## Results

Across 770 trap nights we had 307 human and wildlife detections in the Mount Rainier alpine. We identified at least 16 vertebrate species (seven birds, nine mammals) using glacier-associated habitats (Table 1). Aside from a single camera failure that was unrelated to the trap itself, our low-cost, alpine camera trap design performed well. Across our data set, humans were the most frequently observed taxon (*N* = 208), followed by American pipits (*N =* 18), mountain goats (*N =* 13), hoary marmot (*N* = 13), and wolverines (*N* = 8; Table 1, Figure 4). Just under one-fifth of birds could not be identified to species but for those that were identified, American pipits and white-tailed ptarmigan accounted for more than half of the observations. However, a number of nondescript brown and gray “blurs” were also recorded, of which many were likely small birds (i.e. gray-crowned rosy finch, horned lark), and thus our results for birds are likely underestimations.

**Table 1.**
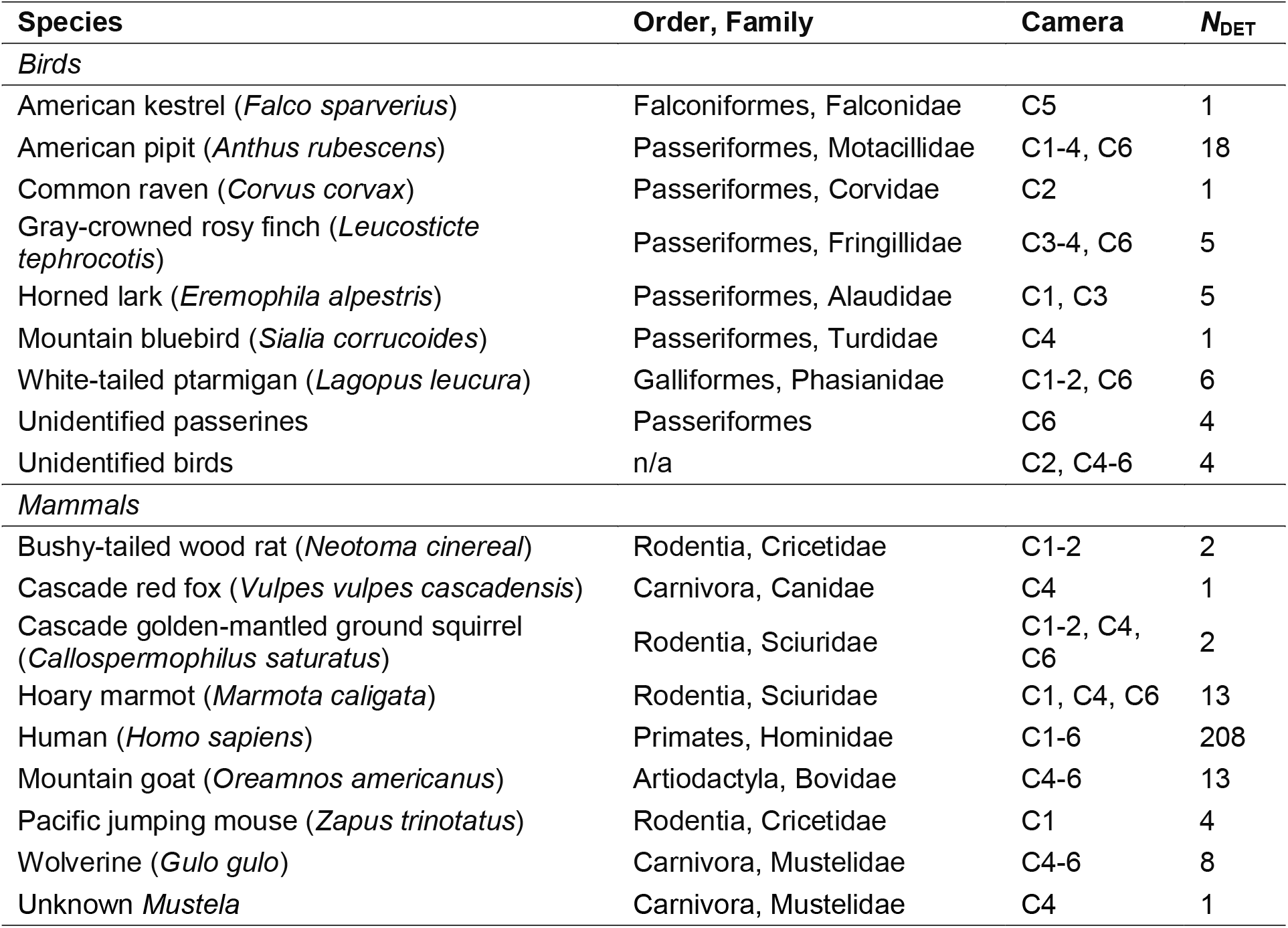
Taxa detected on the Paradise Glacier and directly adjacent habitat of Mount Rainier, WA, USA. *N*_DET_: number of detections. Taxa were classified to the lowest taxonomic possible.

**Figure 4.**
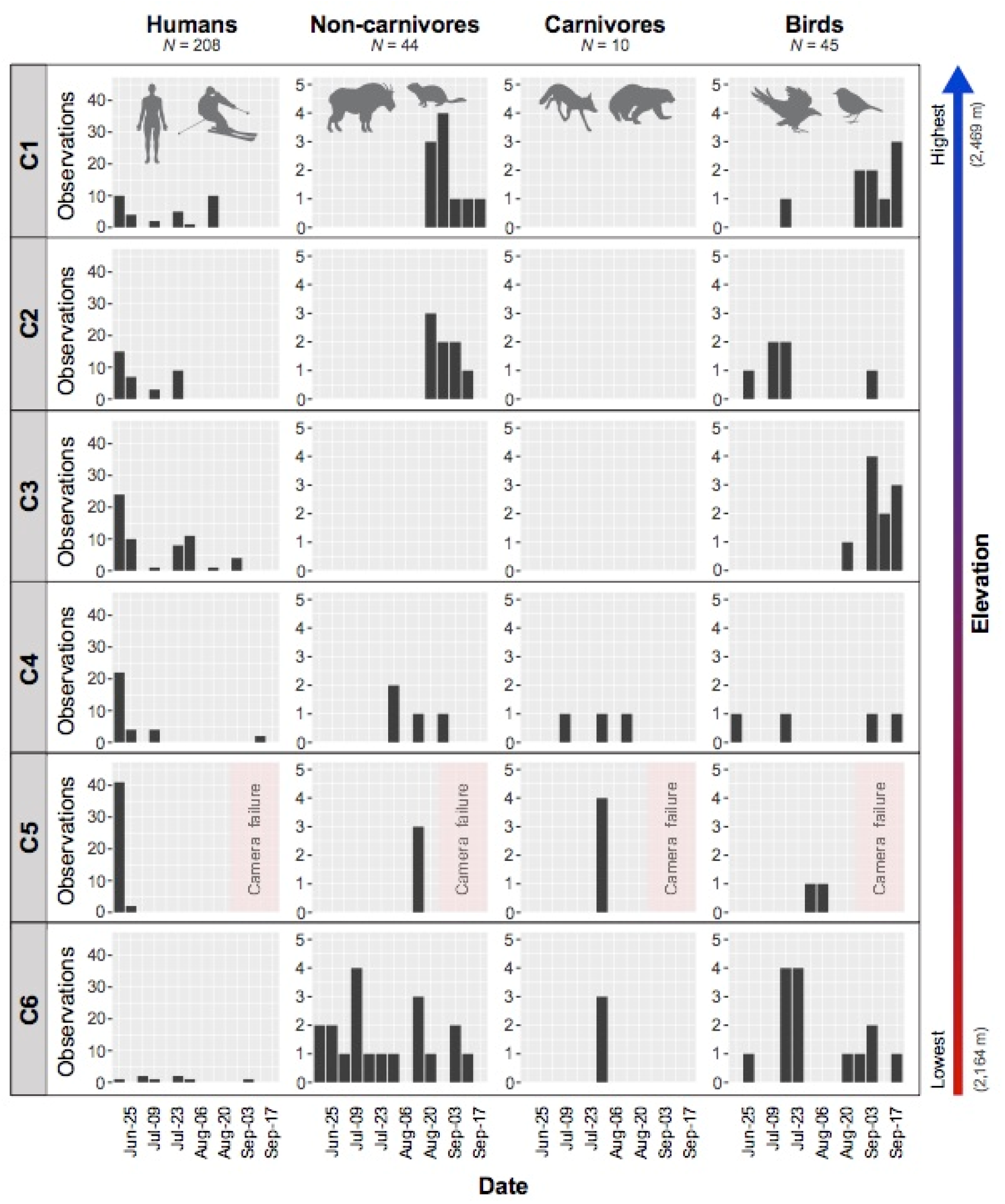
Humans and wildlife observed in this study by elevation (top to bottom) and seasonal timing (grouped on a per week basis; left to right for each taxonomic group). Observations refer to unique sightings and the total observations for a given taxonomic group are given at the top of the plot (N_ALL_). The viewer should note the substantial scale difference of the y-axis for human observations relative to wildlife. A short-term camera failure at C5 is denoted by the labeled light pink square.

We observed the most humans at the third-highest camera, C3 (*N* = 59), and the most wildlife at the lowest camera, C6 (*N* = 36; Figure 4). Interestingly, C6 also recorded the fewest humans (*N* = 8) and the most taxonomically diverse wildlife: five mammals (golden-mantled ground squirrel, hoary marmot, human, mountain goat, wolverine), three identifiable birds (American pipit, gray-crowned rosy finch, white-tailed ptarmigan), and several unidentifiable passerines and other birds. Because we did not control for the amount of area each camera could “see,” comparisons across cameras (and detection periods since cameras were adjusted throughout the study), should be interpreted with caution.

We observed the highest number of detections during the first week of the study (18-24 June 2021) but this pattern was overwhelmingly driven by humans (Figure 4). Indeed, ∼54% of all human observations were within this period. For wildlife (mammals and birds), no clear temporal pattern was present. At cameras where humans and wildlife were regularly observed (e.g., C4, C5), we observed a temporal separation with humans being seen early in the season from mid-June to mid-July and other taxa appearing later (after mid-July; Figure 4). However, we did not observe a correlation—positive or negative—between the presence of humans and wildlife across all cameras and weeks of the study (*P*, Spearman’s rho = 0.106).

During the study, we identified three taxa of conservation concern in Washington state: wolverines (including two kits playing on the glacier), a Cascade red fox, and white-tailed ptarmigan (Figures 3e-g). We observed wolverines at the three lowest cameras (C4-C6) in late July/early August, a Cascade red fox at our C4 camera in mid-August, and white-tailed ptarmigans at the two highest cameras (C1-C2) and the lowest camera (C6), including a pair of ptarmigans together in mid-July at C2.

## Discussion

Mountain glaciers are in rapid decline around the world (Hugonnet *et al*. 2021). Concurrent with this recession of glacier ice is the loss of critical habitat for glacier-associated biodiversity (Stibal *et al*. 2020). To date, most of the research emphasis in glacier biology has centered on understanding the distribution and functional roles of microbial life (Hotaling *et al*. 2021). This is not surprising since microbes are the dominant life form in glacier environments and play critical roles in the cryosphere, even altering the albedo of ice and impacting its rate of melt (Ganey *et al*. 2017; Hotaling *et al*. 2021). However, overlooked in this emphasis have been larger vertebrates that use glaciers and adjacent habitats for travel, nesting, foraging, and other key aspects of their life histories (Rosvold 2016). This lack of research emphasis stems, in part, from the difficulties associated with studying wildlife in rugged, glaciated ecosystems, including the limitations of traditional methods when working in harsh, snow-covered habitat.

In this study, we deployed an array of novel camera traps along the margin of a rapidly receding mountain glacier to test our camera design and gain a better understanding of human and wildlife use of these habitats. We found considerable success with our approach and identified an array of birds and mammals, from songbirds to mesocarnivores, highlighting the ecological complexity and potential importance of glaciers and alpine ecosystems to mountain biodiversity. Indeed, many species observed in this study (e.g., American kestrel) have not been previously linked to glacier habitats, raising questions about the degree to which components of their life history have been overlooked. We also found limited, albeit non-significant, evidence for interactions between human and wildlife usage of glacier ecosystems with little to no overlap between the two groups in space and time.

From a conservation and management perspective, we observed three wildlife species that are particularly notable: wolverine (*Gulo gulo*), Cascade red fox (*Vulpes vulpes cascadensis*), and white-tailed ptarmigan (*Lagopus leucura*). All three are listed as “Species of Greatest Conservation Need” under the Washington State Wildlife Action Plan (Washington Department of Fish and Wildlife 2021), with a need for monitoring and distribution information specifically highlighted for wolverine and Cascade red fox. The successful identification of all three species during the course of this study shows the value of wildlife camera trapping above treeline. Furthermore, wolverines are exceptionally rare in the South Cascades with the family (adult female, two kits) observed in this study likely representing just the third family in the range in a century (National Park Service 2021). Given their rarity in MORA and the South Cascades, we assume that all of our observations were of the same wolverines, which were seen on three separate cameras during the week of 30 July 2021. This burst of activity might indicate successful foraging in the area as wolverines are known to use rendezvous sites near food caches, particularly when kits are young (Inman *et al*. 2012). Notably, we only observed mesocarnivores (e.g., wolverines) where we also observed common prey species (e.g., marmots; Table 1).

While our camera trap design and array in the Mount Rainier alpine was successful, it was not without difficulties. Indeed, the challenging nature of camera trapping in a high-alpine landscape should not be overlooked. In addition to highly variable, and often difficult, weather conditions, the landscape does not lend itself well to traditional camera trapping approaches for three reasons. (1) The physical landscape dramatically changes during summer due to snowmelt. During 2021, seasonal snow depth at a substantially lower elevation than our study area peaked in late winter at 5.36 m (213 inches; Paradise SNOTEL #679). This seasonal snow completely melted by mid-July, effectively lowering the habitat surface by several meters during the first month of our study (mid-June to mid-July). Thus, camera trap locations were limited either to areas that were snow-free early in the melt season or required revisiting as the snow surface changed. In our case, portability was important as we moved cameras small distances throughout the season to track shifting ice margins. Changing snow depth also meant that framing the view, and controlling for the field of vision across cameras, was nearly impossible. For instance, one view that was completely snow-covered in early July was entirely rock and talus within a matter of days. The end of the season offers another complication as seasonal snow can accumulate rapidly, particularly in “sheltered” areas that are ideal for obscuring camera traps. Thus, we recommend that similar future studies in the PNW or comparable areas conclude by the end of September or that researchers devise a strategy for finding snow-covered cameras and/or keeping them free of snow. (2) Hiking and/or game trails are minimal or non-existent above treeline. Thus, an exceptionally large and consistent survey area exists with little to no mechanism for targeting areas where wildlife may be more abundant. To this end, a denser camera array would have certainly yielded better quantifications of vertebrate presence and interactions (e.g., links between carnivores and prey or humans and wildlife). Finally, (3) we did not observe many songbirds on the glacier and in surrounding habitats despite well-known connections between the two (e.g., Hotaling *et al*. 2020). This was likely due to a combination of our cameras not being sensitive enough to detect small birds and images with potential birds in-flight being too blurry for identification. Indeed, a number of photographs included unidentifiable brown or gray blurs that we could not confidently categorize as a bird or something else (e.g., a flying insect). One way to overcome this, or at least improve the detection rate of small birds, would be to pair cameras with acoustic recorders at each site so song could be used in combination with (or in lieu of) imagery.

## Conclusion

In this study, we showed that a simple, cost-effective camera trap design could reveal a rich community of birds and mammals, including species of conservation concern, are using difficult-to-study glacier habitat in western North America. Given the relatively small number of camera traps placed in the vast alpine habitat of Mount Rainier with no game-trails and few natural chokepoints to increase success, it is likely that our results are a significant underestimation of the true scale of biodiversity in these habitats. For instance, in a global meta-analysis, Rosvold (2016) identified 36 species of birds and mammals that are known to use snow and ice habitats for some portion of their life history. Here, we observed 16 species on a small glacier during a single summer with a limited array. Thus, our current understanding clearly underestimates the true scale of wildlife associations with mountain glaciers. As glaciers continue to rapidly recede worldwide, there is a growing need to document and understand these wildlife-ice connections to properly account for the biodiversity impacts of cryosphere decline.

## Acknowledgements

This research was conducted under Mount Rainier National Park permit #MORA-2021-SCI-0016. We thank Ben Lee for assistance in the field and park staff for their support during the permitting process. We also thank David Sousa and the co-owners of Tatoosh Meadows for allowing us to camp on their property during this research. We gratefully acknowledge that our research was conducted on the traditional lands of the Cowlitz, Muckleshoot, Nisqually, Puyallup, Squaxin Island, and Yakama tribes.

## Data availability

The data and scripts used in this study are available in this study’s GitHub accession: https://github.com/scotthotaling/glacier_cams.

